# Cholera toxin-induced disease generates epithelial cell-derived L-lactate that promotes *Vibrio cholerae* growth in the small intestine

**DOI:** 10.1101/2025.08.12.669941

**Authors:** Maria de la Paz Gutierrez, Jenna L. Hall, Sebastian E. Winter, Fabian Rivera-Chávez

**Affiliations:** Division of Host-Microbe system and Therapeutics, Department of Pediatrics, University of California San Diego, 9500 Gilman Dr; La Jolla, CA 92093, USA; Department of Molecular Biology, School of Biological Sciences, University of California San Diego, 9500 Gilman Dr; La Jolla, CA 92093; Division of Infectious Diseases, School of Medicine, University of California at Davis, One Shields Ave; Davis CA 95616, USA

## Abstract

Cholera toxin (CT) promotes *Vibrio cholerae* colonization by altering gut metabolism to favor pathogen growth. We have previously found that CT-induced disease leads to increased concentrations of L-lactate in the lumen of the small intestine during experimental cholera. Here, we show that CT-induced disease leads to the upregulation of mammalian lactate dehydrogenase A (LDHA), an enzyme that catalyzes the conversion of pyruvate to L-lactate, in small intestinal epithelial cells. In a suckling mouse model, the bacterial L-lactate dehydrogenase (LldD) conferred a fitness advantage to *V. cholerae* but not to the Δ*ctxAB* mutant incapable of producing CT. Finally, the fitness advantage conferred by LldD was significantly reduced in mice lacking epithelial-cell specific LDHA, demonstrating that epithelial-derived L-lactate is a major contributor to CT-dependent pathogen expansion. These findings identify L-lactate as a host-derived metabolite generated by intestinal epithelial cells produced during cholera disease that directly fuels *V. cholerae* growth during infection, uncovering a mechanism by which CT confers a fitness advantage to the pathogen during disease.

## INTRODUCTION

*Vibrio cholerae* is the causative agent of cholera, a severe secretory diarrheal disease that remains endemic in many parts of the world. Following ingestion, *V. cholerae* colonizes the small intestine, where it produces cholera toxin (CT), a potent enterotoxin that activates host adenylate cyclase and drives fluid secretion into the intestinal lumen (Faruque et al., 1998). This secretory response results in the profuse watery diarrhea known as “rice water stool,” which underlies both disease symptoms and fecal-oral transmission of the pathogen.

In addition to promoting fluid loss, CT alters the metabolic landscape of the gastrointestinal tract. During infection, CT-induced disease drives marked changes in the composition of host-derived metabolites in the intestinal lumen of infant rabbits, including elevated levels of long-chain fatty acids (LCFAs), heme, and L-lactate. These changes coincide with a transcriptional program in *V. cholerae* enriched for genes involved in the tricarboxylic acid (TCA) cycle, iron acquisition, fatty acid metabolism, and L-lactate metabolism (Rivera-Chávez et al., 2019). The pathogen’s acquisition of both LCFAs and heme has been shown to promote growth during CT-induced disease. Together, these findings suggest that CT not only drives diarrheal symptoms but also establishes a nutrient-rich environment that favors pathogen expansion. However, the mechanisms by which CT-dependent metabolic remodeling confers a fitness advantage to *V. cholerae* remain poorly understood.

Lactate is a glycolytic by-product produced under both aerobic and hypoxic conditions. It exists as two stereoisomers, D- and L-lactate, each metabolized by distinct, stereospecific dehydrogenases. While members of the gut microbiota can produce both isomers, mammalian tissues generate only L-lactate, primarily via the activity of LDHA. Several bacterial pathogens, including *Salmonella* Typhimurium and *Campylobacter jejuni*, utilize host-derived L-lactate during colonization of cecum and colon, respectively (Gillis et al., 2018; Sinha et al., 2024). Notably, epithelial-derived L-lactate has been shown to support the bloom of *Enterobacteriaceae* in the lumen of the large intestine (Taylor et al., 2022). Although increased levels of L-lactate have been observed in the small intestinal lumen during CT-induced disease, the source of this metabolite remains undefined. Moreover, it is unknown whether L-lactate contributes to *V. cholerae* growth in the small intestine.

*V. cholerae* encodes a putative L-lactate dehydrogenase (LldD) and an L-lactate permease (LldP), both of which are transcriptionally upregulated in a CT-dependent manner during infection of the terminal ileum of the small intestine (Rivera-Chávez et al., 2019). However, whether *V. cholerae* actively utilizes L-lactate *in vivo* remains unknown. In parallel, the contribution of CT-induced changes in host epithelial metabolism to *V. cholerae* colonization is also not well understood.

Here, we investigated whether CT-induced disease promotes the production of epithelial-derived L-lactate in the small intestine and whether *V. cholerae* utilizes this metabolite *in vivo* to gain a fitness advantage during infection.

## RESULTS

### LldD is required for utilization of L-lactate by *Vibrio cholerae*

We have previously observed that during infection, L-lactate accumulates in the lumen of the small intestine and *V. cholerae* upregulates the expression of *lldD*, the gene encoding the L-lactate dehydrogenase in a CT-dependent manner (Rivera-Chávez et al., 2019). To determine whether L-lactate alone can induce *lldD* expression, we cultured *V. cholerae* in the presence of L-lactate and found a significant increase in *lldD* expression (p < 0.05; Fig. 1A), supporting the idea that *in vivo* induction of *lldD* during CT-induced disease reflects the availability of L-lactate in the intestinal lumen. Next, to investigate the role of L-lactate dehydrogenase in *V. cholerae* growth, we compared the fitness of the *V. cholerae* wild-type (using Δ*lacZ* for blue-white screening) and Δ*lldD* strains during growth with L-lactate. The *V. cholerae* wild-type strain exhibited a significant (p < 0.001) fitness advantage over the *ΔlldD* mutant under aerobic conditions (Fig. 1B), consistent with a role for LldD in aerobic L-lactate metabolism. A similar significant fitness advantage was observed under microaerobic (p < 0.001) and anaerobic (p < 0.05) conditions in the presence of nitrate (Fig. 1B), indicating that LldD supports growth across a range of physiologically relevant oxygen levels and alternative electron acceptors.

**Figure 1.**
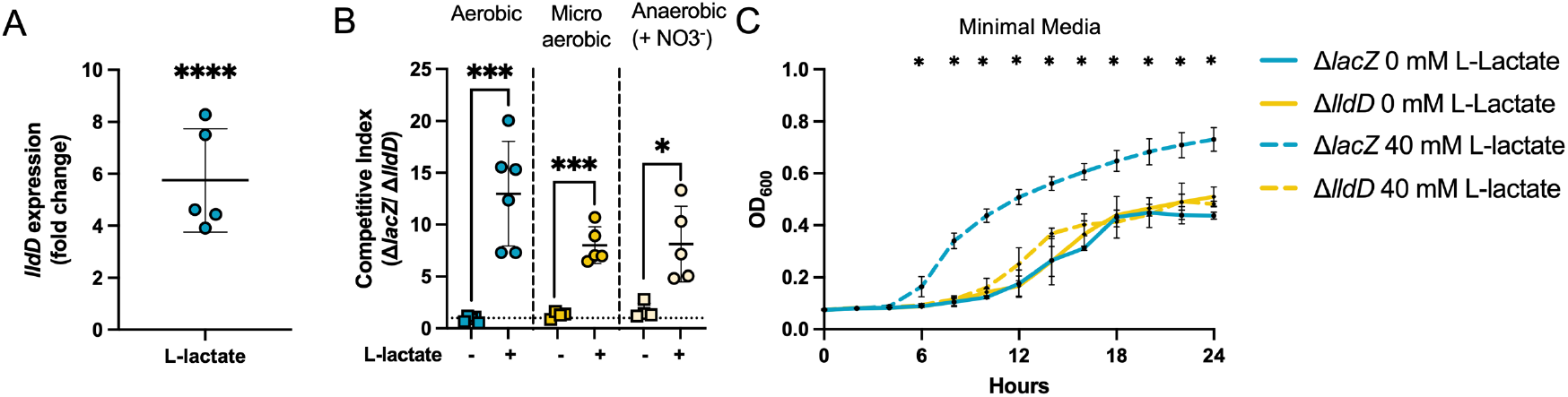
*V. cholerae* LldD is required for L-lactate utilization. (A) Expression of *lldD* was measured on wild-type *V. cholerae* growing in M9 media 0.4% v/v glycerol in the presence or absence (control) of 40 mM L-lactate. Values represent fold change relative to control (n=5). **(B)** Competitive growth assays *in vitro* were performed between a LldD-expressing strain (Δ*lacZ*) and a Δ*lldD* strain in the presence or absence of 40 mM L-lactate. M9 0.4% v/v glycerol media was inoculated with a 1:1 mixture of Δ*lacZ* and Δ*lldD* and incubated for 16 h at 37°C in aerobic, microaerobic and anaerobic conditions. The competitive index (CI) was calculated as the ratio of Δ*lacZ*/Δ*lldD* recovered compared to the ratio of the inoculum (n= 4-6). **(C)** Growth curve of *V. cholerae* Δ*lacZ* (blue) and Δ*lldD* (yellow) in M9 0.4% v/v glycerol media (full line) or M9 0.4% v/v glycerol supplemented with 40 mM L-lactate (dotted line) grown for 24 h at 37°C (n= 4). Data are represented as mean +/-SD. Each dot represents one biological replicate. For statistical analysis, unpaired t-tests against same condition were performed. * p < 0.05; *** p < 0.001; **** p < 0.0001.

In agreement with the competition results, the Δ*lldD* mutant showed significantly (p < 0.05) impaired growth compared to the *V. cholerae* wild-type strain when grown in the presence of L-lactate (Fig. 1C), but not in nutrient rich media (Supplementary Figure 1A). Importantly, LldD also conferred a growth advantage to *V. cholerae* strains incapable of producing CT (i.e., Δ*ctxAB* mutant background) confirming that CT production does not impact L-lactate utilization (Supplementary Figure 1B and Supplementary Figure 1C).

Together, these findings demonstrate that LldD is required for efficient utilization of L-lactate by *V. cholerae* and confers a fitness advantage under diverse environmental conditions that are likely encountered in the gut environment where oxygen levels are low and nitrate becomes available (Bueno et al., 2018).

### L-lactate supports the luminal growth of *V. cholerae* in the small intestine

Since mammalian cells only generate the L-isomer of lactate through the activity of LDHA and we had previously found that CT-induced disease leads to elevated levels of L-lactate in the small intestinal lumen of infant rabbits (Rivera-Chávez et al., 2019), we next asked whether CT-induced disease has the potential to influence the availability of host-derived L-lactate through the direct upregulation of the mammalian L-lactate dehydrogenase gene (*Ldha*). To this end, expression of the host *Ldha* was quantified in the small intestine. *Ldha* expression was significantly elevated (p < 0.05) in both the distal and proximal small intestine in animals infected with wild-type *V. cholerae* but remained low in animals infected with the Δ*ctxAB* strain (Fig. 2A and Fig. 2B). Furthermore, Δ*ctxAB*-infected mice that had been simultaneously treated orally with purified CT had significantly (p < 0.01) elevated *Ldha* levels similar to those observed in wild-type-infected mice (Fig. 2A and Fig. 2B), demonstrating that CT is required to induce expression of host *Ldha*, which would be predicted to lead to elevated L-lactate production in the lumen.

**Figure 2.**
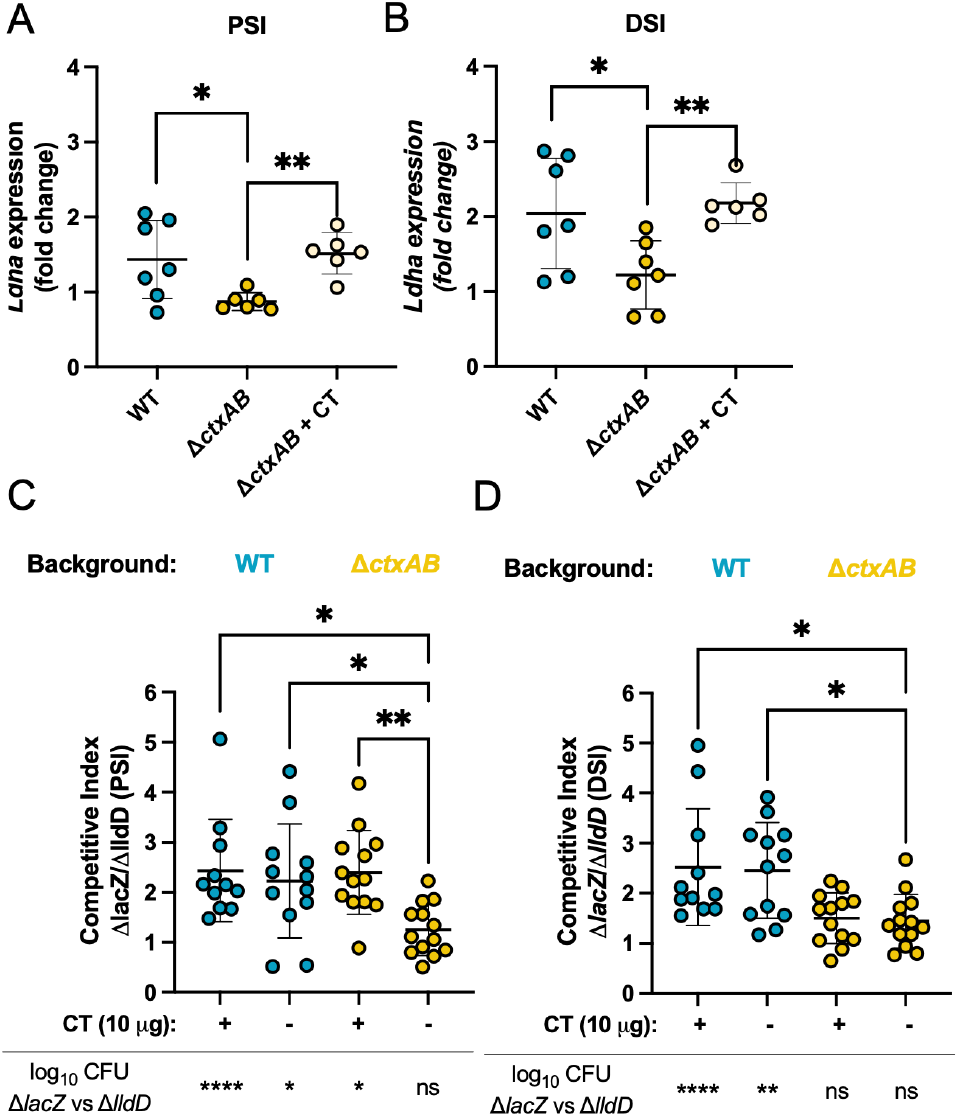
CT-dependent L-lactate utilization confers a fitness advantage to *V. cholerae* in the small intestine. Expression of host L-lactate dehydrogenase *Ldha* was measured in the PSI **(A)** and DSI **(B)** of 7-day-old mice infected with either wild-type (WT), Δ*ctxAB*, or Δ*ctxAB* co-inoculated with CT (10 μg). Mock-infected mice were used as control. Values represent fold change relative to mock-infected control (n= 6-7). Statistical significance was assessed using a one-way ANOVA followed by a Tukey’s multiple comparison test. Competitive assays in the suckling mouse model were performed using a LldD-expressing strain (Δ*lacZ*) and a Δ*lldD* strain. 7-day-old pups were infected with a 1:1 mixture of Δ*lacZ* and Δ*lldD* with same genetic background: WT (blue) of Δ*ctxAB* (yellow). Bacteria recovered 22 h after infection from the PSI **(C)** and DSI **(D)** were used to calculate CI. When indicated, inoculums were supplemented with purified CT (10 μg) (n=11-13). Data are represented as mean +/-SD. Each dot represents data from one animal. For statistical analysis a Krustal-Wallis test followed by a Dunn’s multiple comparison test was performed. Paired t-test were performed when comparing log_10_ CFU. * p < 0.05; ** p < 0.01; **** p < 0.0001; ns, not statistically significant.

To assess whether L-lactate supports the growth of *V. cholerae* in the lumen of the gastrointestinal tract during infection, 7-day-old suckling mice were infected with a 1:1 mixture of the *V. cholerae* wild-type and the Δ*lldD* mutant (CT-producing strains; left side, blue dots). After 22 hours of infection, the wild-type *V. cholerae* significantly outcompeted the Δ*lldD* mutant in the lumen of the proximal (p < 0.05) and distal (p < 0.01) small intestine, indicating that L-lactate utilization contributes to *V. cholerae* growth *in vivo* (Fig. 2C and Fig. 2D).

Since CT-induced disease leads to increased L-lactate concentrations in the lumen of the small intestine of infant rabbits and upregulation of L-lactate utilization genes in the pathogen (Rivera-Chávez et al., 2019), we next asked whether this L-lactate-dependent fitness advantage required CT-induced disease. To test this, we repeated the competitive infection using a Δ*ctxAB* mutant background which is unable to produce CT (right side, yellow dots). Consistent with the hypothesis that CT is required for L-lactate production, the competitive advantage conferred by LldD was significantly reduced (p < 0.05) in the absence of CT and no fitness advantage was observed between Δ*ctxAB* and Δ*ctxAB ΔlldD* strains, suggesting that L-lactate-dependent growth requires CT-induced disease (Fig. 2C and Fig. 2D). In wild-type infected mice, concurrent exogenous CT treatment did not significantly increase the competitive advantage of the Δ*lacZ* strain over the Δ*lldD* mutant, consistent with the idea that CT produced during wild-type infection is sufficient to drive maximal L-lactate–mediated growth. Importantly, infection with the Δ*ctxAB* and Δ*ctxAB ΔlldD* strains concurrently with oral CT-treatment led to a significant (p < 0.05) fitness advantage for the Δ*ctxAB* mutant in the proximal small intestine, further supporting that CT-induced disease is responsible for L-lactate availability to enable pathogen growth in the small intestine (Fig. 2C).

Together, these findings demonstrate that CT promotes the generation of host-derived L-lactate in the small intestine and that *V. cholerae* benefits from the availability of this metabolite through LldD-dependent utilization of L-lactate during infection.

### Small intestinal epithelial cells are the primary source of L-lactate during CT-induced disease

The intestinal epithelium is the primary target of CT (Heyningen et al., 1976), which acts on epithelial cells in the small intestine to drive ion secretion and water loss during infection (Faruque et al., 1998). We next asked whether small intestinal epithelial cells are also the source of the L-lactate produced during CT-induced disease. Treatment of IEC-6 cells, a rat small intestinal epithelial cell line, with purified CT led to a significant (p < 0.05) increase in *Ldha* expression, supporting a role for small intestinal epithelial cells in L-lactate production during infection (Fig. 3A). *Ldha* induction in IEC-6 cells was significantly (p < 0.05) abrogated by inhibition mechanisms by which it enhances of mTORC1 signaling with rapamycin, consistent with a model in which CT stimulates *Ldha* expression via mTORC1-dependent activation of HIF-1α (Kierans et al., 2020).

**Figure 3.**
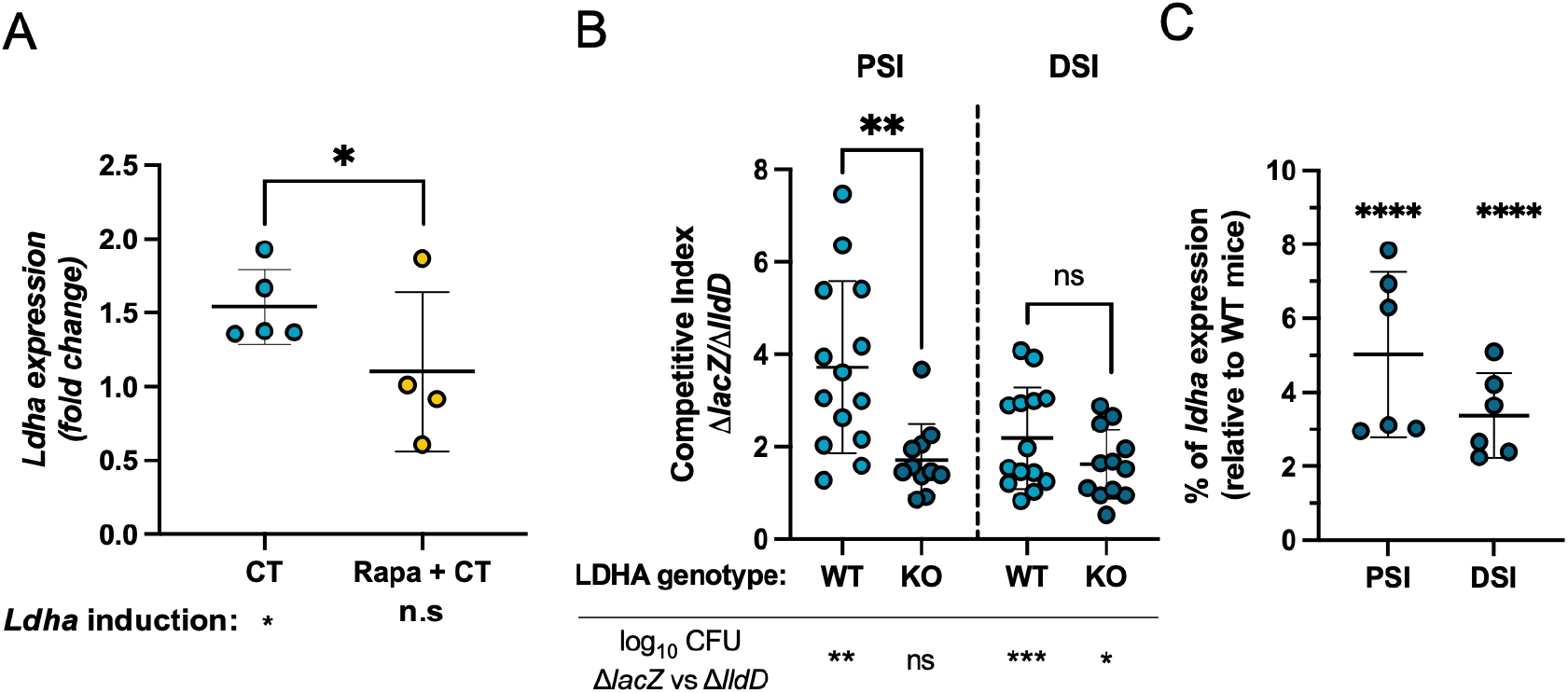
Epithelial cell–derived L-lactate supports *V. cholerae* growth in the small intestine. (A) Expression of L-lactate dehydrogenase *Ldha* was measured in IEC6 cells 2 h post-CT (10 μg) treatment. Where indicated, cells were pre-treated with 200 nM rapamycin for 18 h before the addition of CT. For both treatments, media was used as control (n=4-5). **(B)** Competitive assays between a LldD-expressing strain (Δ*lacZ*) and a Δ*lldD* strain were conducted in 7-day-old LdhaΔIEC mice and wild-type (WT) littermates (n=11-14). **(C)** Expression of *Ldha* in LdhaΔIEC mice infected with *V. cholerae*, shown as percentage of *ldha* expression in WT mice infected with *V. cholerae* (n=6). Data are represented as mean +/-SD. Each dot represents data from one animal or one well. Unpaired t-test were performed for statistical analysis of CI and *Ldha* expression levels compared between treatments (A) or to WT-infected mice (C). Paired t-test were performed when comparing log_10_ CFU. * p < 0.05; ** p < 0.01; *** p < 0.001; **** p < 0.0001; ns, not statistically significant.

To examine whether epithelial-derived L-lactate contributes *V. cholerae* growth in the small intestine, we infected epithelial-specific *Ldha* knockout mice (*Ldha*^ΔIEC^ mice) and littermate controls with a 1:1 mixture of the *V. cholerae* wild-type and the Δ*lldD* mutant. Consistent with our previous observations that L-lactate metabolism supports luminal growth in the small intestine (Fig. 2C and Fig. 2D), the *V. cholerae* wild-type strain (Δ*lacZ*) significantly (p < 0.05) outcompeted the Δ*lldD* mutant in littermate controls (Fig. 3B). Importantly, this fitness advantage was significantly reduced in the proximal small intestine of *Ldha*^ΔIEC^ mice, indicating that *V. cholerae* primarily relies on epithelial-derived L-lactate to support growth in the lumen of the proximal small intestine during infection. Further supporting this conclusion, *Ldha* transcript levels were reduced by over 90% (p < 0.0001) in both proximal and distal small intestines of *Ldha*^ΔIEC^ mice compared to littermate controls infected with wild-type *V. cholerae*, confirming that epithelial-specific deletion effectively abrogates CT-dependent *Ldha* induction during infection (Fig. 3C).

These findings identify epithelial cells in the small intestine as the primary source of CT-induced L-lactate and demonstrate that this host-derived metabolite promotes *V. cholerae* growth during infection. Together, these results uncover a novel mechanism by which CT, a phage-derived bacterial toxin, reprograms host epithelial metabolism to generate a nutrient that *V. cholerae* has evolved to exploit for growth in the gut during disease. This work has broad implications for the function of other bacterial toxins and advances our understanding of how toxin-mediated diarrheal disease confers a fitness advantage to enteric pathogens.

## DISCUSSION

Many pathogenic bacteria secrete toxins that elevate intracellular cyclic AMP (cAMP) or other cyclic nucleotides, leading to widespread disruption of host signaling and tissue function. While the disease-causing effects of these toxins have been well studied, recent work has begun to reveal how toxins can also reshape host metabolism to promote bacterial growth. Although CT was first described over six decades ago (De et al., 1959), the mechanisms by which it enhances *V. cholerae* fitness during infection have remained unclear.

Here, we identify a novel function for CT in the remodeling of host epithelial metabolism to generate a nutrient that directly fuels pathogen growth. We show that CT induces the expression of LDHA in epithelial cells of the small intestine, leading to increased availability of L-lactate in the lumen of the small intestine. *V. cholerae* exploits this metabolite using its own L-lactate dehydrogenase (LldD), which enables efficient use of L-lactate as carbon source and respiratory electron donor in the small intestine during disease. To our knowledge, CT is the first virulence factor identified to increase luminal L-lactate during infection, as no such single factor has been identified in infections caused by other pathogens.

Mammalian cells exclusively generate the L-isomer of lactate via LDHA, while gut microbes can produce both L- and D-lactate. *V. cholerae* genetically encodes the ability to import L-lactate via LldP and oxidize it using LldD. While *lldP* and *lldD* are strongly induced in the presence of CT during infection, their functional roles *in vivo* had not been previously defined. Here, we demonstrate that LldD confers a CT-dependent fitness advantage during infection, and that this advantage is significantly diminished in the small intestine of mice lacking LDHA specifically in intestinal epithelial cells. These findings establish epithelial cells in the small intestine – the site where the pathogen colonizes and produces the toxin – as a key source of L-lactate during CT-induced disease. Notably, the modest residual advantage of wild-type *V. cholerae* in LDHA-deficient mice suggests that additional sources of L-lactate – potentially from non-epithelial cells such as phagocytes

– may also contribute to pathogen growth.

The ability of enteric pathogens such as *Salmonella* Typhimurium to manipulate the gut environment and reach high luminal densities is essential for fecal-oral transmission (Lawley et al., 2008). Our findings support the emerging paradigm that diarrheal disease functions not merely as a pathological consequence of infection, but as an evolved strategy to enhance pathogen growth and transmission. CT promotes massive fluid loss, and the resulting “rice water stool” can contain more than 10^11^ *V. cholerae* per liter. These results uncover how CT-induced changes in host metabolism promote pathogen expansion in the gut, offering new insight into how toxin activity may contribute to fecal-oral transmission.

More broadly, this work reveals a direct, evolutionarily selected link between a horizontally acquired phage-encoded toxin, host metabolism, and bacterial nutrient acquisition. It raises the possibility that other bacterial toxins may operate through similar metabolic mechanisms. For example, pertussis toxin, which shares structural and enzymatic similarities with CT, may reprogram host metabolism in analogous ways during *Bordetella pertussis* infection. In sum, these findings expand our understanding of how microbial toxins contribute to pathogen ecology and evolution by modulating host physiology to promote pathogen growth and transmission.

## METHODS

### Bacterial strains and growth conditions

*V. cholerae* and *Escherichia coli* strains used in this study are listed in Supplementary Table S1. *V. cholerae* and *E. coli* were routinely grown aerobically at 37°C in LB broth or on LB plates. When indicated, media was supplemented with streptomycin 100 μg/mL, carbenicillin 100 μg/mL, chloramphenicol 5 μg/mL (*V. cholerae*), chloramphenicol 20 μg/mL (*E. coli*) or X-gal. For infections, *V. cholerae* was grown in LB broth in the absence of antibiotics for 16 h.

For RNA extraction, wild-type *V. cholerae* was grown aerobically for 20 h in M9 media supplemented with 0.4% v/v glycerol and streptomycin in the presence or absence of 40 mM sodium L-lactate. After the incubation, bacteria were pelleted, resuspended in 1 ml of TRIzol reagent (Invitrogen, cat# 15596018) and stored at -80°C until used.

### Generation of *V. cholerae* mutants

To generate a suicide plasmid for the clean deletion of *ctxAB*, a fragment containing two segments of homologous DNA flanking the genomic region of *ctxAB* (ctxABF1F2) was obtained by digesting pDS132::ctxABF1F2 with KpnI and XbaI (NEB). The released fragment was cloned into the plasmid pRE107 digested with the same enzymes. The resulting plasmid pRE107::ctxABF1F2 was used for allelic replacement in *V. cholerae*.

Similarly, to generate a suicide plasmid for the clean deletion of *lldD* two segments of homologous DNA flanking the genomic region of *lldD* (lldDF1F2) were obtained by PCR and cloned into the plasmid pDS132 using Gibson Assembly. The resulting plasmid pDS132::lldDF1F2 was used for allelic replacement in *V. cholerae*.

The suicide plasmids were introduced into *V. cholerae* by conjugation with *E. coli* SM10. Transconjugants were selected on LB plates supplemented with streptomycin and carbenicillin (for integration of pRE107::ctxABF1F2) or streptomycin and chloramphenicol (for integration of pDS132::lldDF1F2). Transconjugants were cultured overnight in LB (low sodium) in the absence of antibiotics and then plated onto LB with 15% w/v sucrose to select clones that had lost the plasmid. Mutants were confirmed by PCR using primers flanking the deleted region. All primers used are listed on Table S2.

### Growth curve *in vitro*

*V. cholerae* grown on a LB plate was used to inoculate a M9 liquid culture supplemented with 0.4% v/v glycerol and streptomycin and grown for 6 h on a shaker at 37°C. The liquid culture was then diluted in M9 media to an OD_600_ of 0.22 and subsequently diluted 1/10^th^ in the same media with or without 40 mM sodium L-lactate (Sigma-Aldrich, cat# L7022). Two-hundred microliters of the resulting bacterial suspension were seeded in 96-well plate and incubated at 37°C with shaking in a plate reader SpectraMax ID5 (Molecular Devices) that measure OD_600_ every 30 min for a total of 24 h.

### Competitive growth experiments *in vitro*

*V. cholerae* Δ*lacZ*, Δ*lldD*, Δ*ctxAB* Δ*lacZ* and Δ*ctxAB* Δ*lldD* grown on a LB culture overnight were diluted in M9 media supplemented with 0.4% v/v glycerol and streptomycin to an OD_600_ of 0.22 (equivalent to 2×10^8^ CFU/mL). The bacterial suspensions containing the Δ*lacZ* and the Δ*lldD* strains (or Δ*ctxAB* Δ*lacZ* and Δ*ctxAB* Δ*lldD*) were mixed in a 1:1 ratio and diluted to 2×10^6^ CFU/mL in the same media. For competitive growth experiments in aerobic and microaerobic condition, 5 mL of M9 0.4% v/v glycerol 100 μg/mL streptomycin with or without 40 mM L-lactate were inoculated with 50 μl of the inoculation mix (1×10^4^ CFUs). For experiments in anaerobic conditions cultures were additionally supplemented with 20 mM sodium nitrate. Cultures were incubated for 16 hours at 37°C in a shaker (aerobic condition), in static with the lid completely closed (microaerobic condition) or in an anaerobic chamber (AnaeroPack, MGC) The inoculums as well as the cultures after incubation were serial diluted in PBS and plated in LB plates supplemented with streptomycin and X-gal. Competitive indeces were calculated as: (Output Δ*lacZ* / Output Δ*lldD*) / (Input Δ*lacZ* / Input Δ*lldD*).

### Suckling mouse model

7-day-old C57BL/6 pups were separated from their dams 1-hour prior infection. Pups were randomly allocated to experimental groups and orally gavaged with 1×10^6^ CFU *V. cholerae* alone or with 10 μg of CT in 50 μl of LB and placed inside a 30°C incubator. For competitive infections, *V. cholerae* strains were mixed in a 1:1 ratio and diluted to 1×10^6^ in 50 ul. Alternatively, pups were given LB as mock infection control. After 22 h, pups were euthanized, and the GI tracts were aseptically removed. The small intestine was cut into two pieces (proximal and distal), and the contents were squeezed and collected in 1 mL of PBS each. Tissues were snapfrozen in liquid nitrogen and stored at -80°C until used. Bacterial burden was determined by plating serial dilutions of the contents onto LB plates, supplemented with streptomycin. For competitive *in vivo* experiments, both inoculum and contents were plated onto plates containing streptomycin and X-gal. Competitive indeces were calculated as (Output Δ*lacZ* / Output Δ*lldD*) / (Input Δ*lacZ* / Input Δ*lldD*).

All experiments in this study were approved by the Institutional Animal Care and Use Committee at the University of California San Diego.

### Cell culture

IEC6 cells were grown in Dulbecco’s Modified Eagle’s Medium (DMEM, Corning, cat# 10013CV) supplemented with 10% v/v fetal bovine serum (FBS) and 1% v/v penicillin-streptomycin in a 5% CO atmosphere at 37°C. Twelve-well plates were seeded with 2×10^5^ per well and the next day, media was replaced with fresh media alone or containing 200 nM rapamycin (Sigma-Aldrich, cat# 553210). After 18 h, CT was added to the wells in a final concentration of 100 μg/ml and the cells were incubated at 37°C for 2 h. After the incubation, media was removed and 1 ml of TRIzol reagent was added to each well to lyse the cells. Samples were stored at -80°C until RNA was extracted.

### RNA extraction and qRT-PCR

For RNA extraction, 200 μl of chloroform were added to bacterial or mammalian cells samples (resuspended in 1 ml of TRIzol reagent). Alternatively, snapfrozen proximal and distal small intestine samples were added to Green RINO RNA lysis kit (Next Advance) containing 1 ml of TRIzol reagent and homogenized for 5 min in Bullet Blender Storm Pro homogenizer (speed 10). The supernatants were added to new tubes containing 200 μl of chloroform. Samples with TRIzol reagent-chloroform were shaken vigorously and incubated for 15 min at room temperature. Samples were centrifuged at 12000 xg for 15 min at 4°C and the aqueous phase was transferred to a new tube. After the addition of 500 μl of 70% ethanol, the samples were transferred into a RNeasy micro column and continued isolation protocol according to manufacturer’s instructions. The concentration of the DNA-free RNA was quantified using Nanodrop One (Thermo Scientific). Samples were diluted to 10 ng/μl and stored at -80°C until used.

For qRT-PCR, SuperScript III Platinum One-Step qRT-PCR master mix kit (Invitrogen, cat# 12574026) was used following manufacturer’s instructions. Two technical replicate reactions with 2 μl of template RNA per sample per target gene were run on a CFX Opus 384 Real-time PCR thermocycler (Bio-Rad). Gene expression was normalized to a housekeeping gene (*actB* or *gyrB*) using the 2^-ΔΔCT^ method.

### Statistical analysis

Statistical analysis was performed using GraphPad Prism version 10. For comparison between two groups, an unpaired t-test was performed for parametric data and a Mann-Whitney test for non-parametric data. For comparison among three or more groups, a one-way ANOVA test followed by a Tuckey’s multiple comparison test was performed for parametric data, and a Kruskal-Wallis test followed by a Dunn’s multiple comparisons test was performed for non-parametric data. Outliers were determined using the ROUT method (Q = 1).

## SUPPLEMENTARY DATA

Supplementary data include Figures S1 and Table S1

## ACKNOWLEDGMENTS

We thank Bradley Meader for construction of the Δ*lldD* mutant. F.R-C is funded by UC San Diego, the Edward Mallinckrodt, Jr. Foundation, and the Hellman Family Foundation.

## AUTHOR CONTRIBUTIONS

F.R.-C., and M.P.G., designed and conceived the study; M.P.G. and J.L.H performed all experiments. S.E.W generated the *Ldha*^ΔIEC^ mice. All authors contributed to data analysis and writing the manuscript.

---

**Table S1.** Bacterial strains used in this study.

| Name | Description | Reference |
| --- | --- | --- |
| *V. cholerae* C6706 | Wild-type El tor C6706 | Lab collection |
| *V. cholerae* $\Delta lacZ$ | Clean deletion of *lacZ* | (Cameron et al., 2008) |
| *V. cholerae* $\Delta lldD$ | Clean deletion of *lldD* | This study |
| *V. cholerae* $\Delta ctxAB$ | Clean deletion of *ctxAB* operon | (Rivera-Chavez et al, 2019) |
| *V. cholerae* $\Delta ctxAB \Delta lldD$ | Clean deletion of both *ctxAB* and *lldD* | This study |
| *V. cholerae* $\Delta ctxAB \Delta lacZ$ | Clean deletion of both *ctxAB* and *lacZ* | This study |
| *E. coli* PDS132::*ctxABF1F2* | SM10 carrying PDS132::*ctxABF1F2* | (Rivera-Chavez et al, 2019) |
| *E. coli* PRE107::*ctxABF1F2* | SM10 carrying PRE107::*ctxABF1F2* | This study |
| *E. coli* PDS132::*lldDF1F2* | SM10 carrying PDS132::*lldDF1F2* | This study |

**Table S2.**
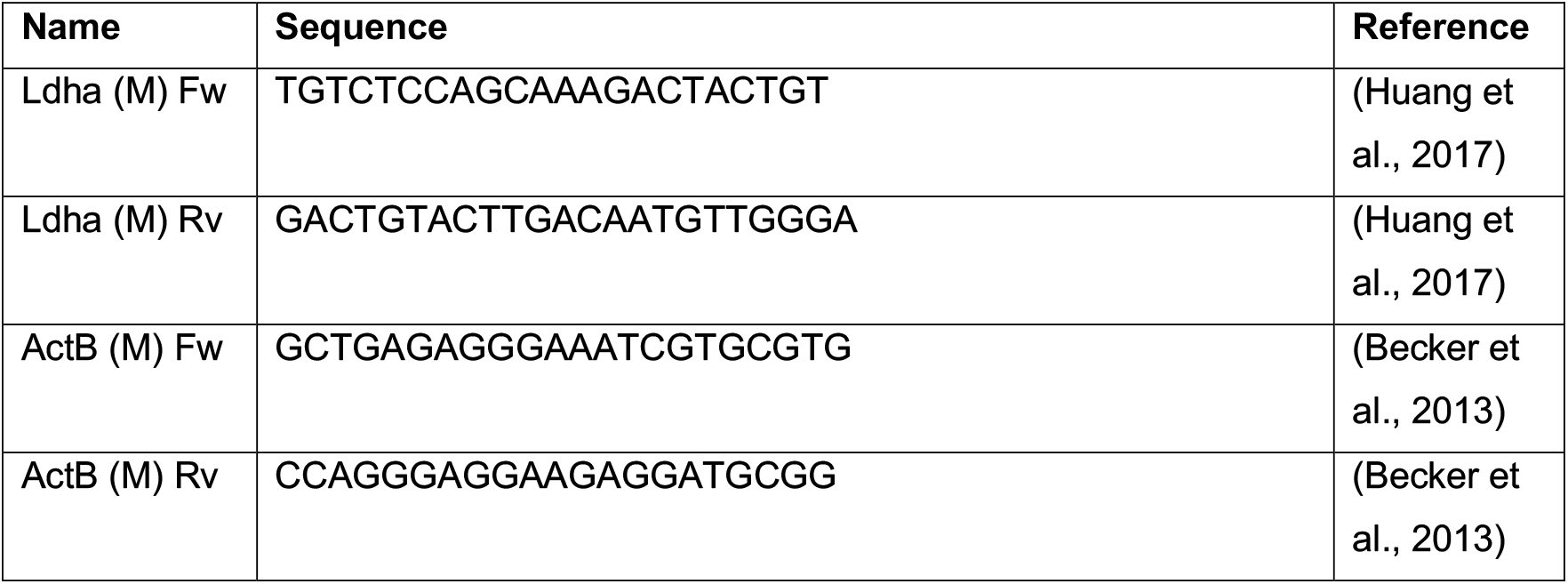
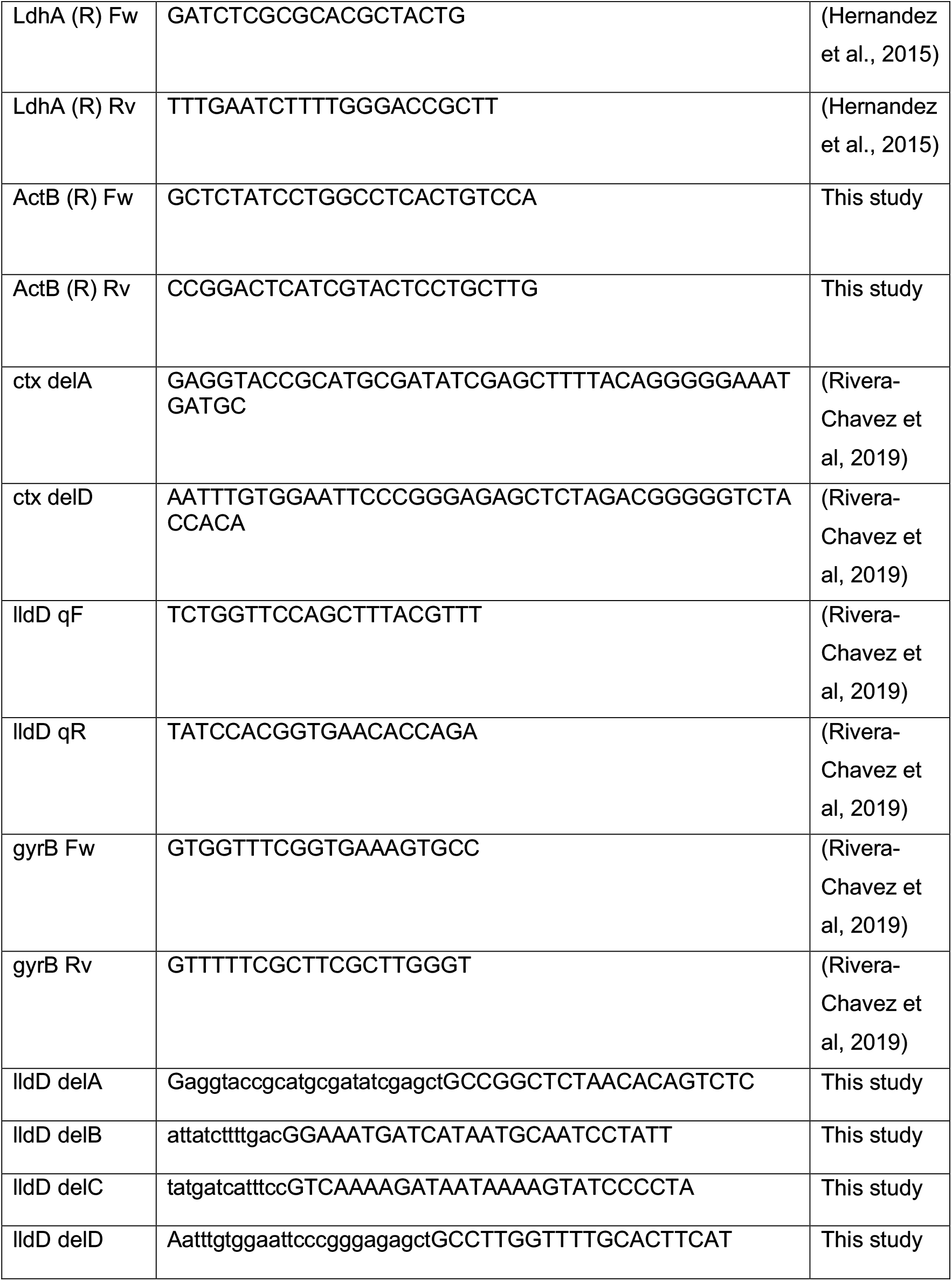
Primers strains used in this study.

**Figure S1.**
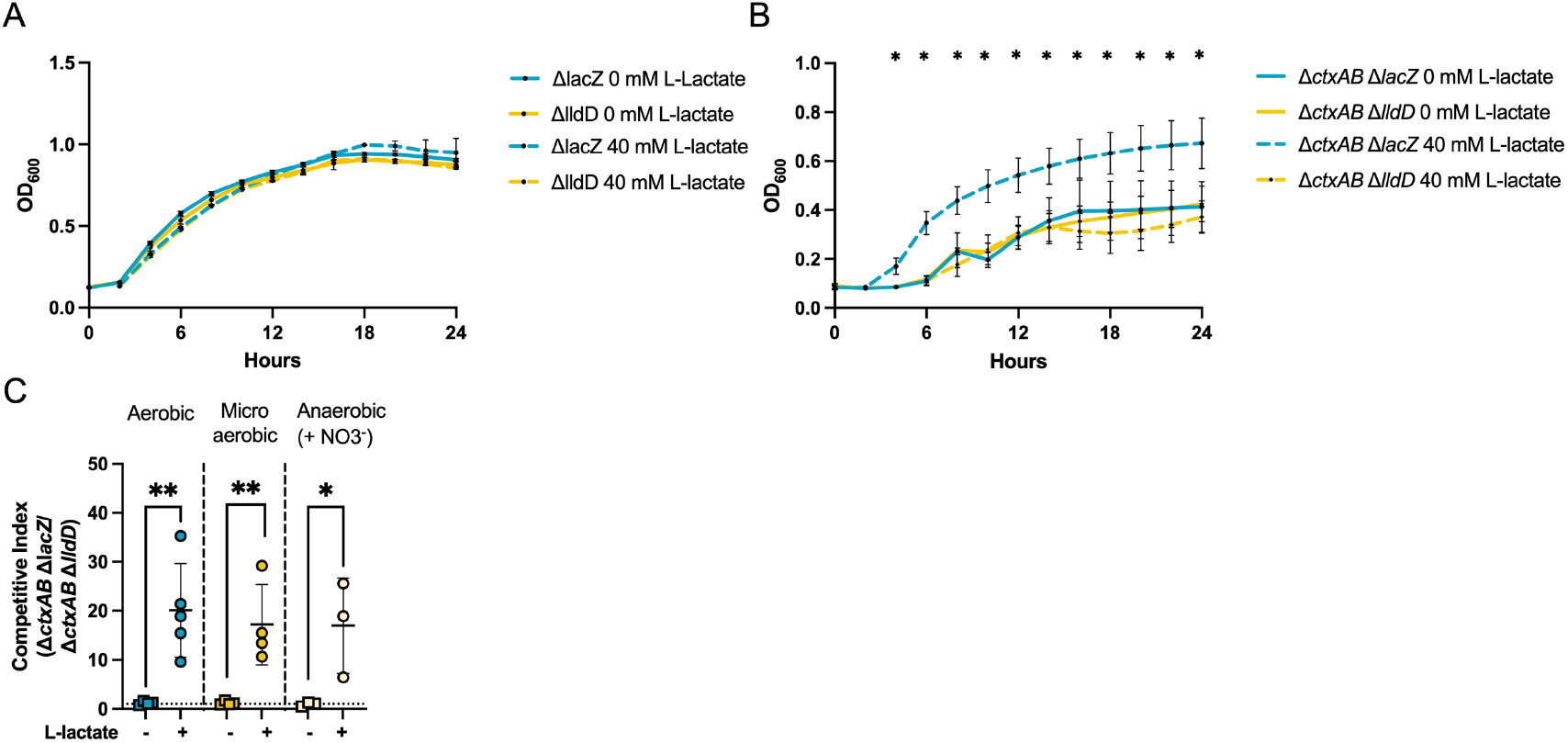
**(A)** Growth curve of *V. cholerae* Δ*lacZ* (blue) and Δ*lldD* (yellow) in LB broth in the presence (dotted line) or absence (full line) of 40 mM L-lactate. **(B)** Growth curve of *V. cholerae* Δ*ctxAB* Δ*lacZ* (blue) and Δ*ctxAB* Δ*lldD* (yellow) in LB broth in the presence (dotted line) or absence (full line) of 40 mM L-lactate. **(C)** Competitive growth assays *in vitro* were performed between Δ*ctxAB* Δ*lacZ* and Δ*ctxAB* Δ*lldD* in the presence or absence of 40 mM L-lactate. The competitive index (CI) was calculated as the ratio of Δ*ctxAB* Δ*lacZ*/Δ*ctxAB* Δ*lldD* recovered compared to the ratio of the inoculum (n= 3-5). Data are represented as mean +/- SD. Each dot represents one biological replicate. For statistical analysis, unpaired t-tests against same condition were performed. * p < 0.05; ** p < 0.01.

## REFERENCES

Becker, S., Oelschlaeger, T.A., Wullaert, A., Pasparakis, M., Wehkamp, J., Stange, E.F., and Gersemann, M. (2013). Bacteria regulate intestinal epithelial cell differentiation factors both in vitro and in vivo. PLoS One 8, e55620.

Cameron, D.E., Urbach, J.M., and Mekalanos, J.J. (2008). A defined transposon mutant library and its use in identifying motility genes in Vibrio cholerae. Proc. Natl. Acad. Sci. U.S.A. 105, 8736–8741.

De, S.N. (1959). Enterotoxicity of bacteria-free culture filtrate of Vibrio cholerae. Nature 183, 1533–1534.

Faruque, S.M., Albert, M.J., and Mekalanos, J.J. (1998). Epidemiology, genetics, and ecology of toxigenic Vibrio cholerae. Microbiol. Mol. Biol. Rev. 62, 1301–1314.

Gillis, C.C., Hughes, E.R., Spiga, L., Winter, M.G., Zhu, W., Furtado de Carvalho, T., Chanin, R.B., Behrendt, C.L., Hooper, L.V., Santos, R.L., et al. (2018). Dysbiosis-associated change in host metabolism generates lactate to support Salmonella growth. Cell Host Microbe 23, 54–64.e6.

Hernandez, A.H., Curi, R., and Salazar, L.A. (2015). Selection of reference genes for expression analyses in liver of rats with impaired glucose metabolism. Int. J. Clin. Exp. Pathol. 8, 3946–3954.

Van Heyningen, W.E., Van Heyningen, S., and King, C.A. (1976). The nature and action of cholera toxin. Ciba Found. Symp. 42, 73–88.

Huang, X., Xie, X., Wang, H., Xiao, X., Yang, L., Tian, Z., Guo, X., Zhang, L., Tang, H., Xie, X., et al. (2017). PDL1 and LDHA act as ceRNAs in triple negative breast cancer by regulating miR-34a. J. Exp. Clin. Cancer Res. 36, 129.

Kierans, S.J., and Taylor, C.T. (2021). Regulation of glycolysis by the hypoxia-inducible factor (HIF): implications for cellular physiology. J. Physiol. 599, 23–37.

Novoa, W.B., Winer, A.D., Glaid, A.J., and Schwert, G.W. (1959). Lactic dehydrogenase. V. Inhibition by oxamate and by oxalate. J. Biol. Chem. 234, 1143–1148.

Rivera-Chávez, F., and Mekalanos, J.J. (2019). Cholera toxin promotes pathogen acquisition of host-derived nutrients. Nature 572, 244–248.

Sinha, R., LeVeque, R.M., Callahan, S.M., Chatterjee, S., Stopnisek, N., Kuipel, M., Johnson, J.G., and DiRita, V.J. (2024). Gut metabolite L-lactate supports Campylobacter jejuni population expansion during acute infection. Proc. Natl. Acad. Sci. U.S.A. 121, e2316540120.

Taylor, S.J., Winter, M.G., Gillis, C.C., Alves da Silva, L., Dobbins, A.L., Muramatsu, M.K., Jimenez, A.G., Chanin, R.B., Spiga, L., Llano, E.M., et al. (2022). Colonocyte-derived lactate promotes E. coli fitness in the context of inflammation-associated gut microbiota dysbiosis. Microbiome 10, 200.

